# Metastatic tumor cells exploit their adhesion repertoire to counteract shear forces during intravascular arrest

**DOI:** 10.1101/443374

**Authors:** Naël Osmani, Gautier Follain, Marìa Jesùs Garcia Leòn, Olivier Lefebvre, Ignacio Busnelli, Annabel Larnicol, Sébastien Harlepp, Jacky G. Goetz

## Abstract

Cancer metastasis is a process whereby a primary tumor spreads to distant organs. We have previously demonstrated that blood flow controls the intravascular arrest of circulating tumor cells (CTCs), through stable adhesion to endothelial cells. We now aim at defining the contribution of cell adhesive potential and at identifying adhesion receptors at play. Early arrest is mediated by the formation of weak adhesion depending on CD44 and integrin αvβ3. Stabilization of this arrest uses integrin α5β1-dependent adhesions with higher adhesion strength, which allows CTCs to stop in vascular regions with lower shear forces. Moreover, blood flow favors luminal deposition of fibronectin on endothelial cells, an integrin α5β1 ligand. Finally, we show that only receptors involved in stable adhesion are required for subsequent extravasation and metastasis. In conclusion, we identified the molecular partners that are sequentially exploited by CTCs to arrest and extravasate in vascular regions with permissive flow regimes.

## INTRODUCTION

Cancer metastasis is a complex multi-step process in which secondary tumors are formed in colonized organs ultimately leading to the death of patients (Lambert et al., 2017). Circulating tumor cells (CTCs) exploit body fluids to reach distant organs, where they will extravasate and either remain dormant or form new tumor foci (Obenauf and Massagué, 2015). The molecular receptors expressed by tumor cells and that are involved in the intravascular adhesion of CTCs have been widely studied (Reymond et al., 2013). In particular, glycoproteins involved in leukocytes rolling at the surface of the endothelium, such as selectins, are required for the adhesion of cancer cells to endothelial cells (Aigner et al., 1998; Laferrière et al., 2001). CD24, CD44, PODXL and mucins facilitate CTC adhesion to the endothelium (Aigner et al., 1998; Dallas et al., 2012; Hanley et al., 2006; Rahn et al., 2005; Shea et al., 2015). Adhesion receptors of the integrin family such as integrins αvβ3, β1 and β4 were similarly involved in adhesion of CTCs to endothelial cells (Felding-Habermann et al., 2001; Laferrière et al., 2004; Klemke et al., 2007; Reymond et al., 2012; Barthel et al., 2013). However, the respective contribution of these receptors in intravascular arrest has rarely been challenged *in vivo* in realistic shear conditions. Furthermore, the correlation between the adhesive and the metastatic potential of CTCs is still not fully understood.

Recently, using a combination of easy-to-tune microfluidics and intravital imaging in two animal models, we have shown that blood flow tunes CTC arrest, preceding metastatic outgrowth, through a tug of war with CTC adhesion potential and thereby control their probability of intravascular arrest (Follain et al., 2018). Using the zebrafish embryo, we observed that CTC arrest immediately upon blood circulation entry but may still be detached from the endothelium by ripping shear forces. Adhesion of CTCs is then reinforced preventing them from shear-mediated detachment. Interestingly, intravascular arrest of leukocytes is driven by weak adhesions with receptors such as CD44, while adhesions reinforcement depends on integrin-mediated bonds (Eibl et al., 2012; McEver and Zhu, 2010).

Here, we demonstrate that highly metastatic CTCs are more prone to intravascular arrest, which favors metastatic extravasation. We show that extravasation is preceded by two major steps: (i) early arrest mediated by the formation of *de novo* adhesions of weak energy (through CD44 and the integrin αvβ3) and (ii) the stabilization of CTC/endothelium bond through the recruitment of adhesions with larger magnitude of energy (via the integrin α5β1). Altogether, these 2 consecutive steps allow arrested CTCs to resist blood flow shear forces and favor metastatic extravasation.

## RESULTS

### CTC adhesive properties correlate with adhesion and extravasation potential

Whether intravascular arrest of CTCs determines their metastatic potential remains to be determined. We thus built on a zebrafish-based experimental metastasis workflow (Follain et al., 2018) and compared the behavior of tumor cells with established low and high metastatic potential derived from a D2 mouse mammary hyperplastic alveolar nodule (Morris et al., 1993). While D20R cells are poorly aggressive, D2A1 cells recapitulate breast carcinoma derived lung metastasis in a BALB/C mouse syngeneic model (Shibue et al., 2013) and were shown to be metastatic in zebrafish embryos (Follain et al., 2018).

When monitoring early arrest of these 2 cell lines, we observed a ~20% decrease in the efficiency of arrest of low-metastatic D20R, when compared to D2A1 (Figure 1A-C). While D2A1 preferentially arrested in the dorsal aorta (DA) and arterio-venous junction (AVJ), D20R were mostly observed in the AVJ and the caudal veins (CV), further along the vascular network (Figure 1C-D). When monitoring the stable intravascular arrest at 3hpi, we confirmed that D20R were mainly found in the CV (Figure 1E-G) suggesting that they have a lower adhesive potential that prevents them from stably arresting in the DA or AVJ. Interestingly, increased arrest in the CV had been observed when flow forces were increased (Follain et al., 2018), confirming the intricate link between shear forces and adhesive potential. We discarded the possibility that increased arrest of D2A1 cells was favored by mechanical trapping of cells with bigger size (i.e. both cell lines display similar size; ~10μm in diameter, Figure S1A). We further checked the surface expression of CD44 and ITGB1, that are respectively weak and strong force adhesion receptors. Interestingly, although expression levels were similar between the two cell lines, surface expression of CD44 and ITGB1 were respectively reduced by ~25% and ~42% in low metastatic D20R cells (Figure S1B-D). Finally, while ~98.5% of D2A1 cells where found extravascular at 24hpi, only ~32% of D20R cells did so (Figure 1H-J, S1E) in vascular regions that match their intravascular arrest location (Figure 1D,F-G,H). Altogether, this demonstrates that the metastatic potential of CTCs is directly linked to their adhesion repertoire, which dictates the efficiency of intravascular arrest.

**Figure 1.**
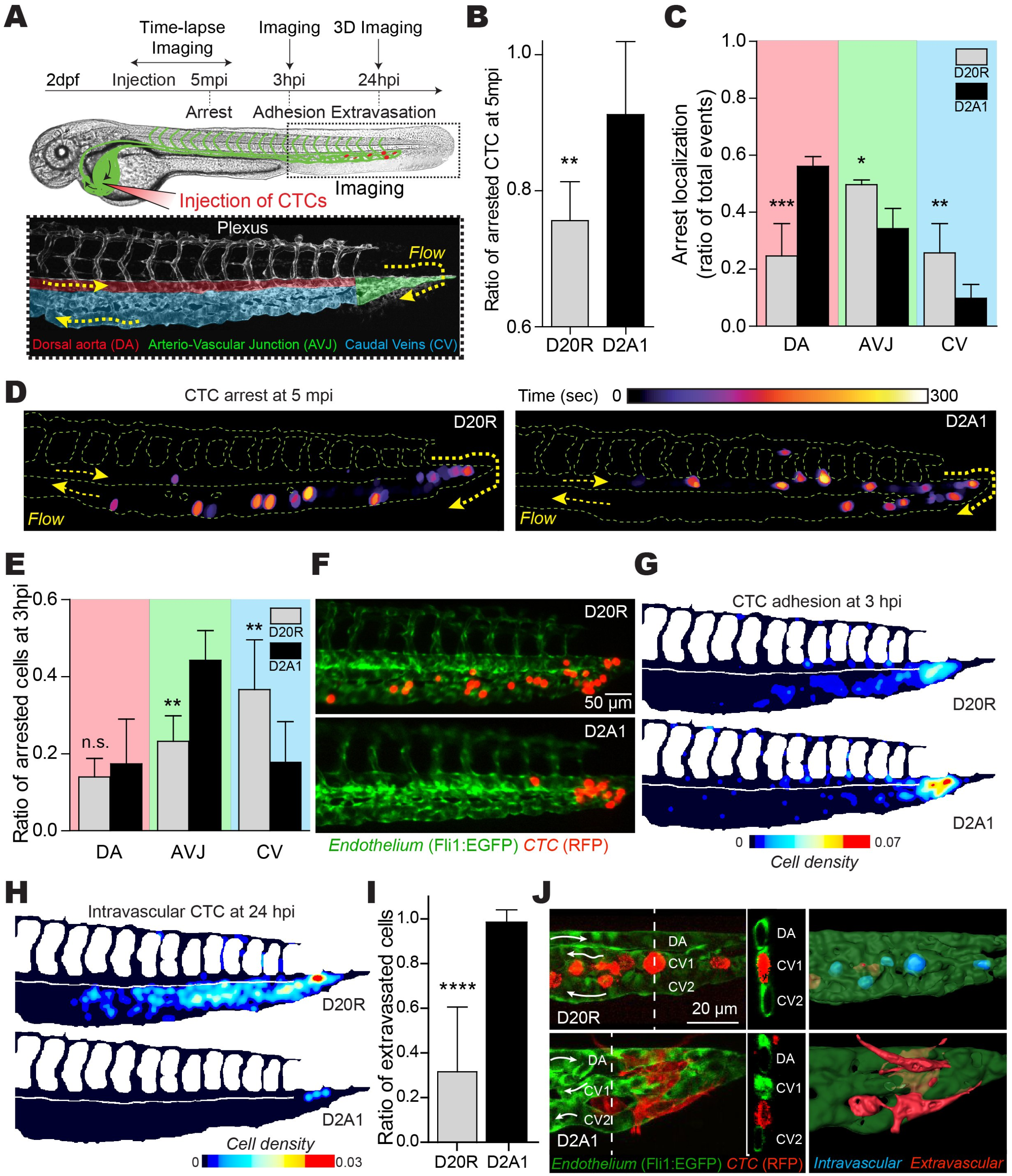
Highly-metastatic D2A1 cells are more prone to stable arrest in larger vessels than D20R cells *in vivo*. (A) Scheme of the experimental approach using 2 dpf *Tg(Fli1:EGFP)* zebrafish embryo as a metastatic cascade model. Lower panel shows blood flow in the plexus regions: dorsal aorta (DA), arterio-venous junction (AVJ) and the caudal veins (CV). (B-D) The arrest of D20R or D2A1 cells were live imaged for 5 min immediately after injection. (B) The ratio of cells stably arrested after 5 min over the total number of cells was measured. (C) The number and localization (see A) of cell arrests over the total number of arrests in the plexus was measured. The graphs show the mean ± S.D. of 3 independent experiments. (D) Time projection of representative embryo injected with indicated cells transfected. The color code shows the arrest time of CTCs. (E-G) The adhesion pattern of D20R or D2A1 cells was imaged 3h after injection. (E) The number and localization (see A) of cells stably adhered at 3hpi was measured. The graph shows the mean ± S.D. of 3 independent experiments. (F) Representative images of cells arrested at 3hpi. (G) The heatmaps show the quantification of the number and location of stably arrested CTCs at 3 hpi in the caudal plexus. (H-I) D20R or D2A1 cells localization pattern was imaged 24h after injection with 3D confocal imaging. (H) The heatmaps show the quantification and location of intravascular CTCs at 24 hpi in the caudal plexus. (I) The ratio of cells extravasated (extravascular) over the total number of cells was measured. The graph shows the mean ± S.D. of 3 independent experiments. (G) Representative images and orthoslice of cells at 24hpi. 3D rendering shows intravascular (blue) and extravascular (red) cells at 24hpi.

### CD44, ITGB3 and ITGB1 favor arrest of CTCs, yet only ITGB1 is required for stable adhesion to the endothelial layer

In order to conduct a molecular dissection of CTC arrest, we built on previous observations which identified a threshold flow value of 400~500 μm/s favoring CTC arrest (Follain et al., 2018). We established a microfluidic approach to probe the CTC adhesion repertoire function in realistic shear flow conditions (Figure 2A-B). Flow-dependent adhesion of CTCs to endothelial cells was probed and challenged by tuning flow velocities as well as adhesion receptors. In control conditions, ~80% of arrested CTCs, whose arrest had been favored by decreasing flow velocities, had formed stable adhesion with the endothelial layer when challenged with increasing flow velocities (Figure 2A-C, Movie 1). Interestingly, while siRNA depletion of CD44, ITGB3 or ITGB1 compromised CTC attachment (Figure 2D, S2, Movie 2-3), only ITGB1 depletion impacted CTC stable adhesion to the endothelial layer (Figure 2D). This suggests that, while CD44 and ITGB3 favor CTC arrest, only ITGB1 allows their stabilization on endothelial cells, in flow conditions, which mirrors their previously described adhesive potential (Bano et al., 2016; Roca-Cusachs et al., 2009).

**Figure 2.**
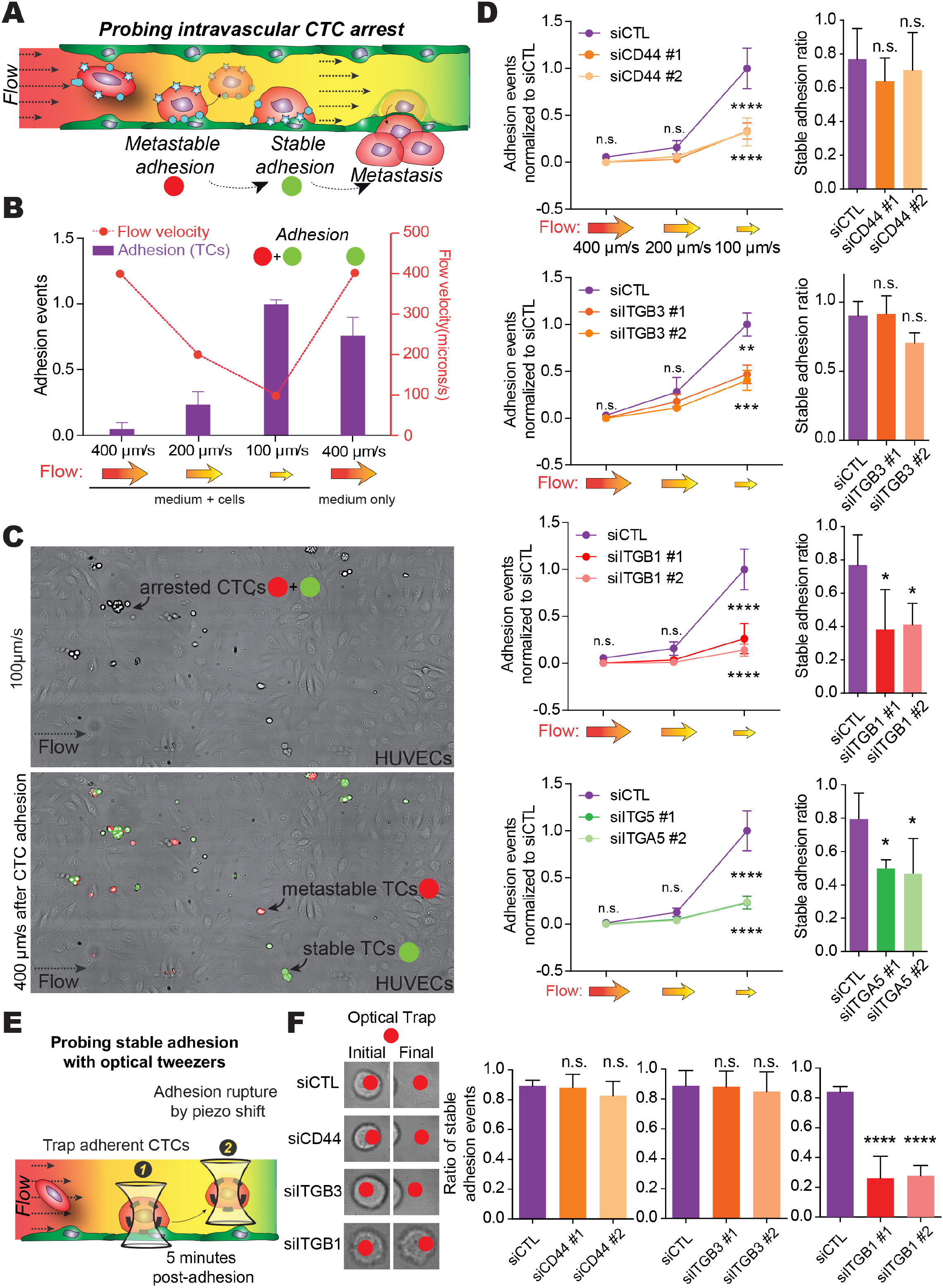
CD44, ITGB3 and ITGB1 are involved in CTC arrest/adhesion but only ITGB1 is required for stable adhesion *in vitro*. (A) Scheme of the experimental approach for the microfluidic CTC arrest assay. (B) D2A1 were perfused into microfluidic channels containing a confluent monolayer of HUVEC cells for 2 min at the indicated speeds. The number of adhered cells is quantified and normalized to the number of cells adherent at the lowest perfusion speed (100 μm/s). The graph shows the mean ± S.D. of 14 independent experiments. (C) Upper panel: time projection of 2 min perfusion of D2A1 cells at 100 μm/s showing arrested cells. Lower panel: time projection of 2 min wash step (perfusion of medium at 400 μm/s) showing transiently arrested cells (metastable, red) or stably arrested cells (stable, green). Related to movie 1. (D) D2A1 cells were transfected with indicated siRNAs and perfused into microfluidic channels containing a confluent monolayer of HUVEC cells. The number of cells adhered normalized to siCTL was quantified (left) and the ratio of stably adhering cells was measured (right). The graphs show the mean ± S.E.M. (left) and mean ± S.D. (right) of 6 independent experiments. Related to movie 2 and 3. (E) Scheme of the experimental approach for the microfluidic CTC stable adhesion assay. (F) D2A1 cells were transfected with indicated siRNAs, perfused into microfluidic channels containing a confluent monolayer of HUVEC cells and left to attach without flow for 5 min. Attached cells were then trapped into the optical tweezer beam and was moved away using the piezo stage. The graph shows the mean ± S.D. of 3 independent experiments. Related to movies 4 to 6.

In order to identify the α subunit (ITGA) mediating stable adhesion of CTCs, and thereby identify its ligand, we depleted ITGA4, 3 and 5 (Figure S2D, S3A). When associated to ITGB1, they respectively mediate binding to luminal VCAM (Klemke et al., 2007), to laminin (Chen et al., 2016a) and to fibronectin (Huveneers et al., 2008) of the basal ECM. We discarded the participation of ITGA4 and observed that ITGA3 was not required for the formation of stable cell adhesion to the endothelial layer (Figure S3B-C). Only ITGA5 depletion phenocopied ITGB1 knockdown and impaired CTC attachment and the formation of stable adhesion (Figure 2D).

In order to specifically discriminate between weak metastable and stable adhesions, we used optical tweezing of adhered CTCs within microfluidic channels. This strategy allowed us to further probe the range of forces that are mediated by CD44, ITGB3 and ITGB1 by assessing the adhesive potential of cells depleted for those receptors. Upon mechanical detachment of CTCs that had established adhesion (Figure 2E), we confirmed that ~80% of control cells formed stable adhesion with the endothelium. Only ITGB1 depletion could impair the number of stable adhesion events (Figure 2F, Movie 4-6). In conclusion, while CD44, ITGB3 and ITGA5/ITGB1 both favor the initial arrest and adhesion of CTCs, only ITGA5/ITGB1 is capable of mediating stable shear-protective adhesion of CTCs to the endothelium.

### CD44 and ITGB3 are required for CTC arrest *in vivo*, only ITGB1 further stabilizes adhesion to the endothelium

To further demonstrate that initial arrest can be decoupled from stable adhesion, we injected D2A1 cells in zebrafish embryo and monitored intravascular arrest using live imaging for 5 minutes (Figure 3A). Depletion of CD44 and ITGB3 impacted early adhesion of CTCs. On the contrary, depleting ITGB1 had no effect and CTCs arrested similarly the AVJ (Figure 3A-B, Movie 7-9). We carefully quantified the exact arrest duration of CTCs, which exceeded 2 minutes in control conditions. CD44- and ITGB3-depleted cells were either unable to arrest or had shorter arrest durations (Figure 3A,C) suggesting decreased adhesive abilities and/or increased sensitivity to shear forces. We further assessed the number and location of stably-adhered CTCs 3hpi and showed that depletion of CD44 or ITGB3 had no effect (Figure 2D-E). On the contrary, ITGB1 depletion drastically reduced the number of cells stably arrested in the AVJ at 3hpi (Figure 2G). Furthermore, while ITGA4 and ITGA3 depletion did not impair CTC adhesion (Figure S3D-E), ITGA5 phenocopied ITGB1-depletion and impaired stable adhesion of CTCs (Figure 3H). This suggests that CD44/ITGB3 are required for the initial arrest while ITGA5/ITGB1 mediate stable arrest *in vivo*.

**Figure 3.**
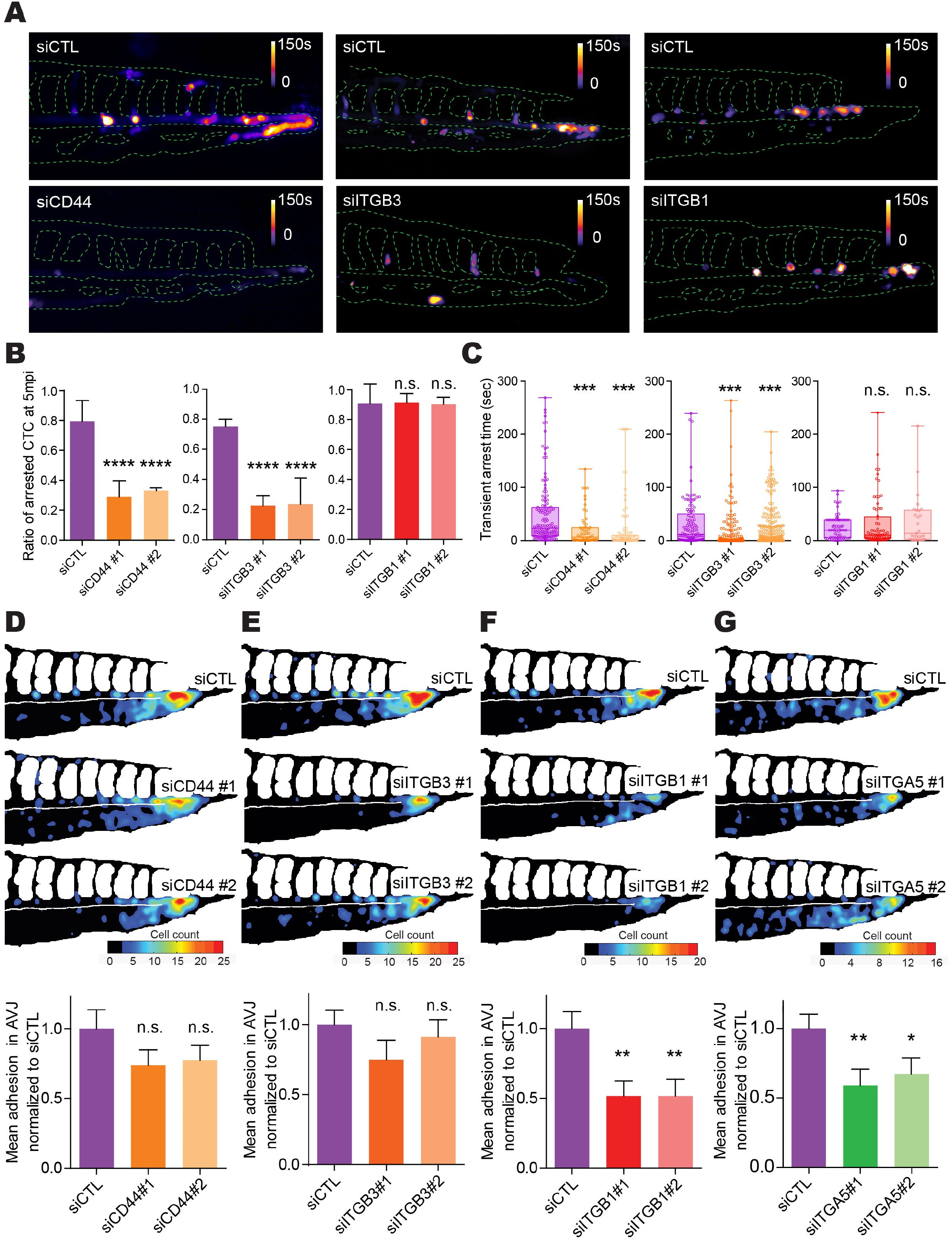
CD44 and ITGB3 are involved in CTC arrest while ITGB1 is required for stable adhesion *in vivo*. D2A1 cells were transfected with indicated siRNAs and microinjected into the duct of Cuvier of 2 dpf *Tg(Fli1:EGFP)* embryos. (A-C) Cell arrests were live imaged for 5 min immediately after injection. (A) Time projection of representative embryo injected with cells transfected with indicated siRNAs. The color code shows the arrest time of CTCs. Related to movies 7 to 9. (B) The ratio of cells stably arrested after 5 min over the total number of cells was measured. The graphs show the mean ± S.D. of 3 independent experiments. (C) Transient arrest time of cells were measured. The graphs show the mean and min/max. of 3 independent experiments. (D-G) Cell adhesion pattern was imaged 3h after injection. The heatmaps show the quantification of the number and location of stably arrested CTCs at 3 hpi in the caudal plexus of embryo injected with indicated siRNAs. The number of cells stably adhered in the AVJ was measured. The graphs show the mean ± S.E.M. of 5 independent experiments.

Fibronectin is a ligand for CD44 and integrins αvβ3 and α5β1 (Jalkanen and Jalkanen, 1992; Humphries, 2006). We next interrogated whether fibronectin could mediate the adhesion-dependent CTC arrest. As expected, we observed flow-dependent fibronectin deposits at the surface of endothelial cells *in vitro* (Figure S4A-B) that are independent of fibronectin expression levels (Figure S4C). Surprisingly, we observed no effect of flow forces on laminin 111 and collagen I, while it decreased both collagen IV and hyaluronan levels (Figure S4D). This suggests that flow forces, that are permissive to CTC arrest (Follain et al., 2018), favor the deposition of fibronectin patches on the luminal side of endothelial cells. We then confirmed such observation *in vivo* and observed luminal fibronectin (Figure S4G). Interestingly, we found that fibronectin was more present in the DA compared to the AVJ thus correlating with higher blood flow velocities (Figure S4F) but also with the early arrest pattern of D21A(Figure 1C-D). This flow dependence was further confirmed as intravascular levels of fibronectin were massively reduced in embryos with reduced blood flow velocities upon lidocaine treatment (Figure S4E,G-H). Altogether, these observations suggest that CTCs synergistically exploit flow forces, flow-mediated deposition of fibronectin and cell surface expression of integrins for efficient intravascular arrest. Doing, they successfully resist to shear forces.

### Efficient intravascular arrest dictates metastatic extravasation

Whether efficient intravascular arrest impact the ability of CTCs to extravasate and grow metastatic colonies remains to be determined. To do so, we focused on CD44 and ITGB1 as mediators of weak and stable adhesion respectively. Interestingly, while CD44 depletion had no effect on metastatic extravasation in zebrafish embryos, ITGB1 depletion significantly reduced the number of cells that underwent extravasation (Figure 4A-C). While such results confirmed the previously identified role for ITGB1 in metastatic extravasation (Stoletov et al., 2010; Chen et al., 2016b), we here provide further insight by demonstrating that ITGB1 favors pro-metastatic intravascular arrest in addition to facilitate transmigration of CTCs (Chen et al., 2016b). To further validate our observation and assess the impact on metastatic outgrowth, we exploited the syngeneic model of lung metastasis upon tail vein injection of luciferase-expressing D2A1 cells in BALB/c mice (Figure 4D). We first carefully assessed seeding efficacy of D2A1 cells using rapid whole-animal imaging of lung signal. Doing so, we noticed that depletion of CD44 only had a mild impact on lung seeding. On the contrary, ITGB1 depletion significantly reduced the number of cells that stably reached the lungs, validating our previous observations (Fig.4E). *Ex vivo* imaging of lungs 13 dpi, which allows to confidently assess the impact on metastatic outgrowth, confirmed that only ITGB1 could impact the metastatic burden (Fig.4F). Altogether, using a unique combination of *in vitro* and two *in vivo* models, we thus demonstrate that stable intravascular arrest of CTC, which is mediated mostly by ITGB1, is a key step preceding successful metastatic outgrowth.

**Figure 4.**
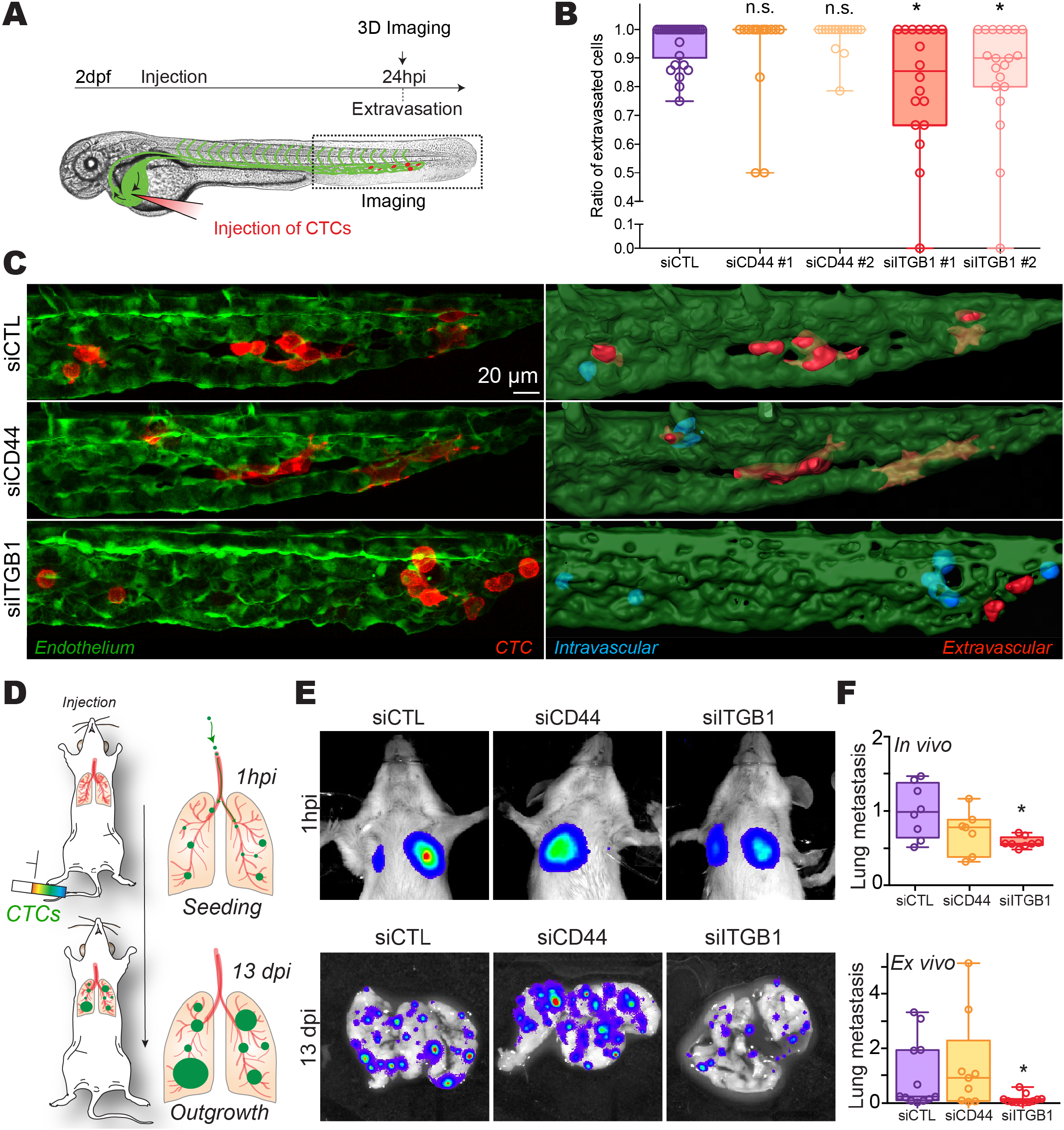
ITGB1 but not CD44 is required for extravasation in the zebrafish embryo and lung metastasis in mice. (A-C) D2A1 cells were transfected with indicated siRNAs and microinjected into the duct of Cuvier of 2 dpf *Tg(Fli1:EGFP)* embryos. Cell localization pattern was imaged 24h after injection. (B) The ratio of cells extravasated (extravascular) over the total number of cells was measured. (C) Representative images of cells at 24hpi. 3D rendering shows intravascular (blue) and extravascular (red) cells at 24hpi. (D) Scheme of the experimental approach for the mouse lung metastasis assay. (E) Representative images of D2A1-Luciferase transfected with the indicated siRNAs in lung seeding (1hpi) and metastatic outgrowth (13 dpi). (F) Quantification of the luciferase signal of D2A1-Luc transfected with the indicated siRNAs in lung seeding (1hpi, upper panel) and metastatic outgrowth (13 dpi, lower panel). The graphs show the mean and min/max. of (B) 5 of or (F) at least 2 independent experiments.

## DISCUSSION

Establishment of metastatic foci follows intravascular journey and efficient arrest of CTCs, the latter being a key step. Although multiple tumor-intrinsic molecular machineries have now been shown to control cancer cell stemness, organotropism, metastatic potential or resistance to chemo/immunotherapy (Lambert et al., 2017), which mechanisms control the stabilization of intravascular arrest, and thereby the efficiency of tumor metastasis, remain to be fully characterized. We provide here the first detailed molecular dissection of this important step by demonstrating how tumor cells exploit their own adhesion machineries, and how they exploit extrinsic factors such as blood flow, to stably arrest in specific vascular regions and favor extravasation, preceding metastatic outgrowth.

Here, we exploited recently-established techniques of intravital imaging of CTCs in the zebrafish embryo, which proves to be a very useful model for such questions (Follain et al., 2018; White et al., 2013). We recently showed that blood flow tunes CTC arrest by a process which is encoded into a tug-of-war between blood flow-driven shear stress and the limited adhesive power of CTCs (Figure 1, Follain et al., 2018). CTC adhesion to the endothelium, preceding their exit from the bloodstream, has been extensively studied either *in vitro* (Aigner et al., 1998; Chen et al., 2016a; Giavazzi et al., 1993; Laferrière et al., 2001, 2004; Reymond et al., 2012; Tremblay et al., 2008) or *in vivo* (Hiratsuka et al., 2011; Köhler et al., 2010; Stoletov et al., 2010) for the past 30 years. These studies identified adhesion receptors used by CTCs to arrest within the bloodstream. However, such molecular dissection, at high spatio-temporal resolution, of CTC arrest and how it dictates metastatic potential *in vivo* remains to be determined. Also, the influence of low energy versus high energy adhesion of CTCs in intravascular arrest *in vivo* remains elusive.

In the present work, we demonstrate that the metastatic potential of CTCs is, in addition to other molecular programs, controlled by their intravascular adhesive potential that dictates their arrest and thereby control their transmigration and metastatic potential. We combined microfluidics and experimental metastasis assays in zebrafish embryos to report that early arrest is mediated by CD44 and integrin αvβ3. Both receptors were reported of weak magnitude (Bano et al., 2016; Roca-Cusachs et al., 2009). Indeed, CD44 has been involved in the rolling of leukocytes along the endothelium (McEver and Zhu, 2010) and is found expressed in breast cancer CTCs with higher metastatic potential (Baccelli et al., 2013; Boral et al., 2017). Integrin αvβ3, a major fibronectin ligand, has been suggested to mediate weaker adhesion involved in mechanosensing during cell migration (Roca-Cusachs et al., 2009; Bharadwaj et al., 2017). We also show that CTC/endothelium bonds are quickly stabilized to resist the shear stress from the blood flow with the formation of integrin α5β1 dependent adhesions. These α5β1-dependent adhesions define more stable adhesions in cells migrating over a matrix of fibronectin as they produce larger magnitude of binding energy but also have longer bond lifetime than integrin αvβ3 (Roca-Cusachs et al., 2009; Kong et al., 2009, 2013; Bharadwaj et al., 2017). These specific biomechanical properties of α5β1 integrins are particularly relevant in a context where CTC adhesion is challenged in a tug of war by shear stress from the blood flow. Finally, we confirm that CTC stable adhesion to the endothelium is mediated by integrin α5β1 both *in vitro* and *in vivo* in the zebrafish embryo. We recently reported that filopodia-like structures are involved in the initial arrest of CTCs (Follain et al., 2018). Interestingly, such structures can carry adhesive receptors such as integrins αvβ3 and α5β1 (Shibue et al., 2012, 2013) suggesting that they might be exploited by CTCs to probe the surface of the endothelium and promote CTC arrest. Importantly, we validated the importance of such step in a lung metastasis assay using a syngeneic mouse model, where we combined fast evaluation of CTC seeding with careful evaluation of the metastatic outgrowth. Doing so, we show that, as expected from the zebrafish experiments, integrin α5β1 controls lung seeding that determines the subsequent metastatic potential in mouse lungs. Altogether, we thus carefully demonstrate that organ seeding and metastatic onset of CTCs are controlled by tumor-intrinsic factors. This is, to our knowledge, the first careful description of how CTCs can exploit permissive blood flow profiles, engage adhesion machineries to efficiently seed metastases.

Recent work suggests that CTCs can engage the laminin in the endothelial basement membrane via integrin α3β1 and α6β1 (Chen et al., 2016b; Wang et al., 2004). We did not observe that integrin α3β1 is required for stable arrest (Figure S3), suggesting that its involvement might be cell line dependent. We did not test the involvement α6β1, and thus cannot rule out its involvement. Finally, we observed that fibronectin, a ligand of CD44 and integrins αvβ3 and α5β1, is found as deposits at the surface of the endothelial layer both *in vitro* and *in vivo*, in a process which is flow-dependent (Figure S4). Fibronectin is found on the luminal side of liver blood vessels and facilitates CTC arrest in this organ (Barbazán et al., 2017). Apical fibronectin secretion by epithelial cells is essential during developmental stages such as cleft formation in mouse (Sakai et al., 2003). Fibronectin is also secreted by endothelial cells, although preferentially basally, and needed for proper vasculogenesis in the zebrafish embryo (Mana et al., 2016). Cellular exocytosis is driven by the increase in membrane tension in response to flow shear stress (Shi et al., 2018). In the zebrafish embryo, we observed a correlation between fibronectin deposits and blood flow profiles (Figure S4F). It is quite tempting to speculate that the shear stress induced by the blood flow might promote the secretion of a fibronectin adhesion platform at the luminal side of the endothelium.

Altogether, we propose that the first steps of CTC arrest are mediated by a coordination between blood flow and CTC/endothelium adhesion where: (i) blood flow shear stress challenges the finite CTC adhesion force to tune their location of arrest, (ii) CTCs differentially use their adhesion repertoire to arrest and stably adhere to the endothelial layer and (iii) only adhesions involved in arrest stabilization are required to further proceed to extravasation and metastatic outgrowth. Targeting such adhesion receptors could be relevant in order to therapeutically prevent CTC extravasation preceding metastatic colonization.

## AUTHOR CONTRIBUTIONS

N.O. & G.F. performed most experiments and analysis. M.J.G.L performed FACS analysis. M.J.G.L and O.L performed and analyzed mice experiments. S.H. supervised the study, performed OT experiments and analysis. J.G. conceived the project and supervised the study. NO wrote the paper with input from G.F., S.H., and J.G.

## ACKNOWLEDGMENTS

We thank all members of the Goetz Lab for helpful discussions. We are grateful to Tsukasa SHIBUE and Robert WEINBERG (MIT) for providing D2A1 cells. We are very grateful to Francesca PERI and Kerstin RICHTER (EMBL) for providing zebrafish embryos. This work has been funded by Plan Cancer (OptoMetaTrap) and CNRS IMAG’IN to S.H. and J.G. and by institutional funds from INSERM and University of Strasbourg. N.O is supported by Plan Cancer (OptoMetaTrap) and the Association pour la Recherche contre le Cancer. G.F. is supported by La Ligue Contre le Cancer and University of Strasbourg. M.J.G.L is supported by INCa (National Cancer institute) and University of Strasbourg (Idex).

## SUPPLEMENTARY FIGURES LEGENDS

**Figure S1.**
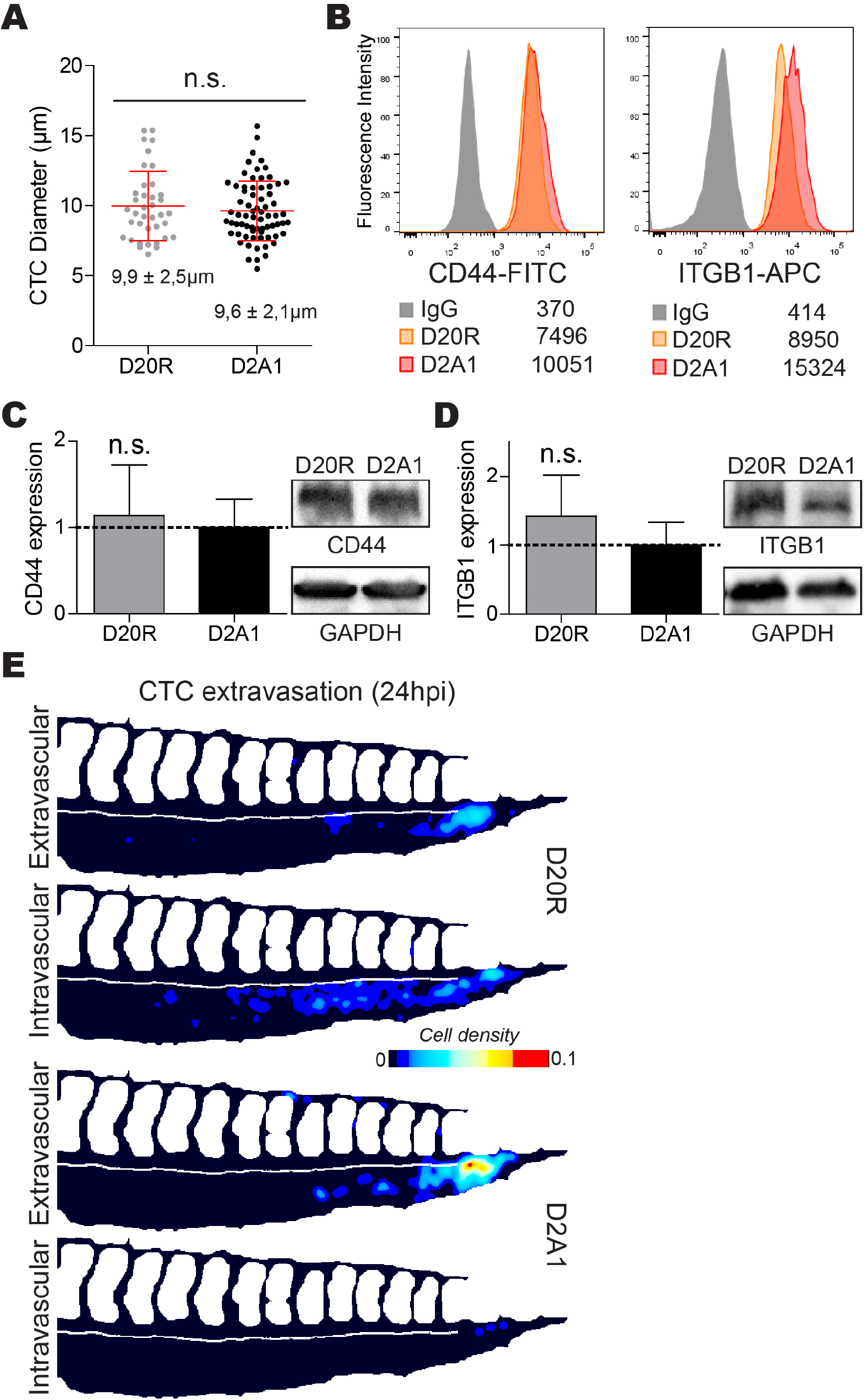
D2A1 and D20R cell line characterization. (A) Quantification of the diameters of D20R and D2A1 cells in suspension. (B) Monoparametric flow cytometry histograms show cell membrane CD44 and ITGB1 levels at the surface of D20R and D2A1 cells after electronic gating on viable cells. FITC- or APC-labelled IgG isotype controls were used in D20R cells to set the negative population. Numbers in legends show the mean fluorescence intensity of the whole populations analyzed. (C-D) Protein extracts were prepared and immunoblotted against (C) CD44 or (D) ITGB1 and GAPDH. The relative expression was measured and normalized to D2A1 expression levels. The graph shows the mean ± S.D. of 5 independent experiments. (E) D20R or D2A1 cells were microinjected into the duct of Cuvier of 2 dpf *Tg(Fli1:EGFP)* embryos and cell localization pattern was imaged 24h after injection with 3D confocal imaging. The heatmaps show the quantification and location of CTCs at 24 hpi in the caudal plexus. Related to figure 1H-J.

**Figure S2.**
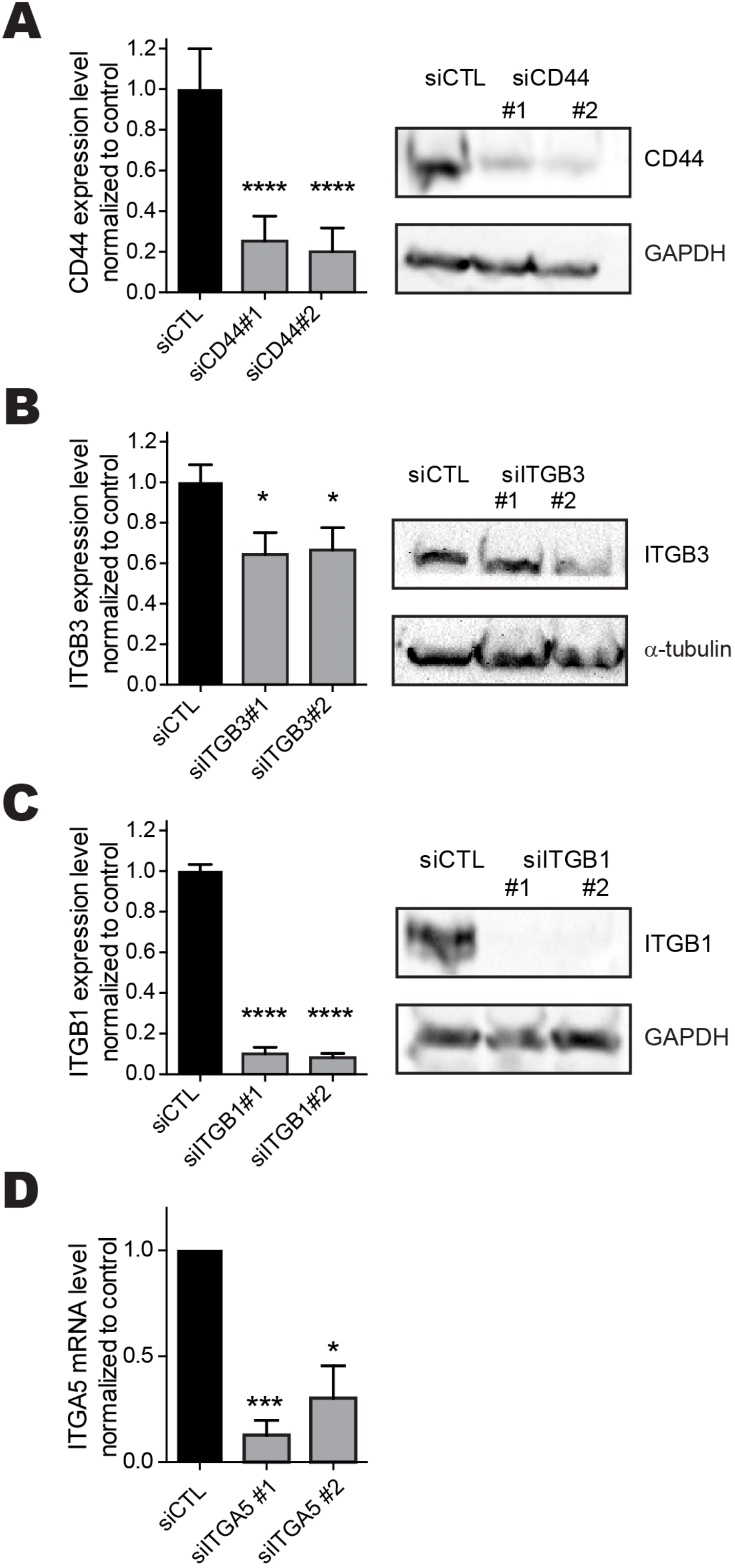
siRNAs knockdown validation after transfection. (A-D) D2A1 cells were transfected with control or (A) anti-CD44, (B) anti-ITGB3, (C) anti-ITGB1 or (D) anti-ITGA5 siRNAs. (A-C) Protein extracts were prepared 72h later and immunoblotted against (A) CD44, (B) ITGB3, (C) ITGB1, (A,C) GAPDH or (B) α-tubulin. The graph shows the mean ± S.D. of 5 independent experiments. (D) Total RNA extracts were prepared 48h after transfection. The expression of ITGA5 was measured using RT-qPCR. The graph shows the mean ± S.D. of 3 independent experiments.

**Figure S3.**
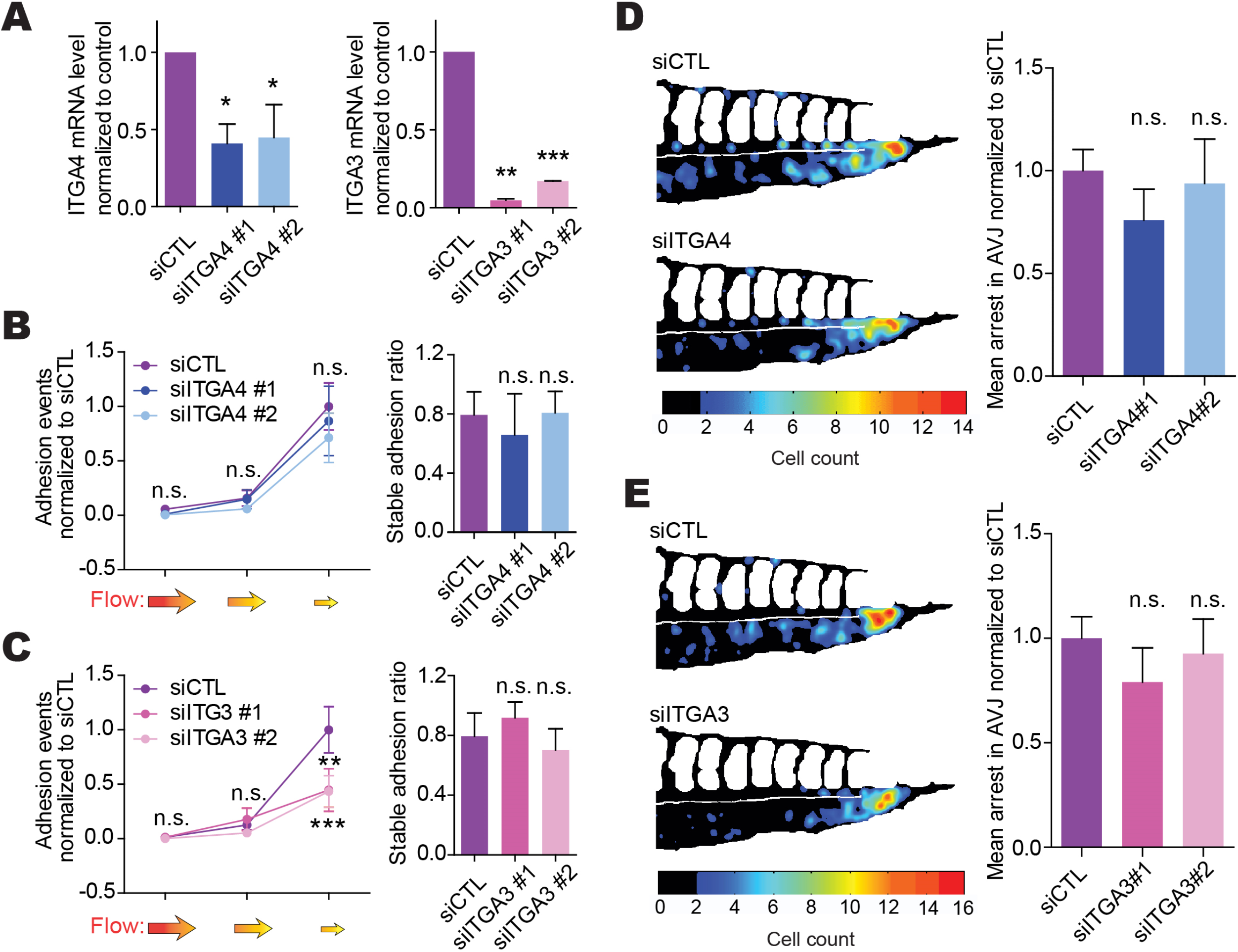
ITGA4 and ITGA3 are not involved in CTC stable adhesion to endothelial cells. (A) D2A1 cells were transfected with indicated siRNAs. The expression of the genes of interest was measured using RT-qPCR. The graph shows the mean ± S.D. of at least 3 independent experiments. (B-C) D2A1 cells were transfected with indicated siRNAs and perfused into microfluidic channels containing a confluent monolayer of endothelial cells (HUVEC). The number of cells adhered normalized to siCTL was quantified (left) and the ratio of stably adhering cells was measured (right). The graph shows the mean ± S.E.M. (left) and mean ± S.D. (right) of at least 4 independent experiments. (D-E) Quantification using heatmapping of the number and location of stably arrested CTCs at 3 hpi in the caudal plexus of embryo injected with cells transfected with indicated siRNAs. The graphs show the mean ± S.E.M. of 4 independent experiments.

**Figure S4.**
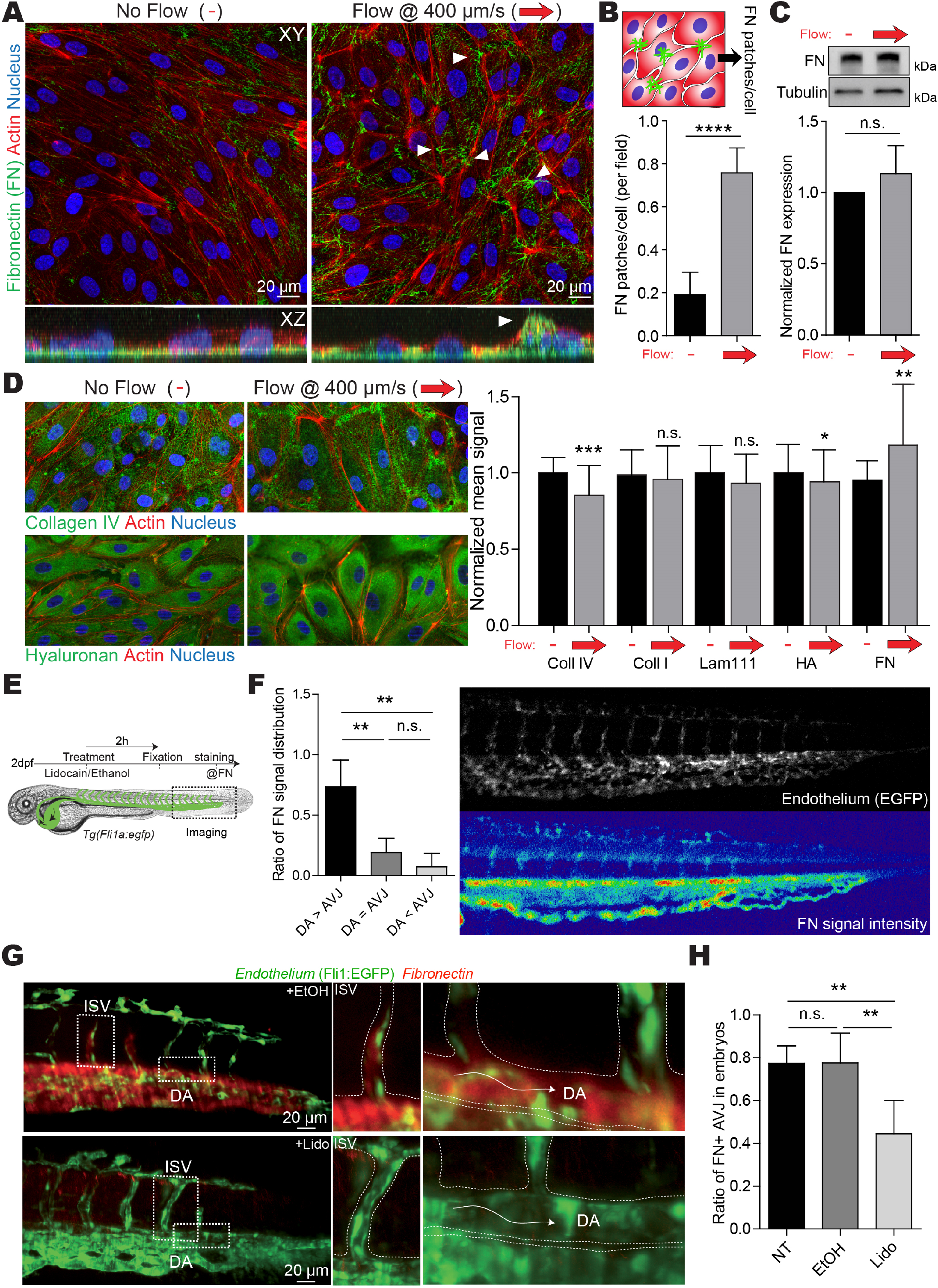
Flow drives the formation of luminal FN deposits. (A) HUVEC cells were grown to confluency in microfluidic channels and subjected to either no flow or a laminar flow of 400 μm/s for 16h. Cells are immunostained for fibronectin (green), actin (red) and nucleus (blue). A y-projection of 35 single representative transversal confocal slices is shown in the bottom panel XZ. (B) The ratio of HUVEC cells with fibronectin (FN) deposits was quantified. The graphs show the mean ± S.D. of 3 independent experiments. (C) HUVEC cells were grown to confluency in microfluidic channels and subjected to either no flow or a laminar flow of 400 μm/s for 16h. Fibronectin (FN) expression was then quantified. A representative western blot image is shown in the upper panel The graphs show the mean ± S.D. of 5 independent experiments. (D) HUVEC cells were grown to confluency in microfluidic channels and subjected to either no flow or a laminar flow of 400 μm/s for 16h. Cells were immunostained for collagen IV, collagen I, laminin 111, hyaluronan (HA) or fibronectin (FN). (right) Representative image of cells stained for collagen IV or hyaluronan (green), actin (red) and nucleus (blue). (left) For each individual field of view, the mean signal of the green channel (ECM) was measured and normalized to the average signal in the no flow condition. The graphs show the mean ± S.D. of 5 independent experiments. (E) Scheme of the experimental approach for the *in vivo* luminal FN quantification. (F) Fibronectin signal in the DA and the AVJ was compared. Embryos were classified as higher signal in the DA (DA>AVJ), identical (DA=AVJ) or higher in the AVJ (DA<AVJ). The graph shows the mean ± S.D. of 6 independent experiments. (G) Representative embryos were imaged using light sheet microscopy (SPIM). (H) 2 dpf *Tg(Fli1:EGFP)* embryos were either untreated (NT) or treated for 2h with vehicle (EtOH) or lidocain at 640 μM (Lido). The presence of fibronectin deposits in the AVJ was assessed. The graph shows the mean ± S.D. of 6 independent experiments.

**Movie 1 – Live adhesion assay**. D2A1 cells were transfected with control siRNAs. 72h later, cells were detached and perfused into microfluidic channels for 2 min at the indicated speeds. The movie is acquired at 24 fps and displayed at 72 fps. Related to Figure 2B-C.

**Movie 2 – Live adhesion movie of siCTL, siCD44 and siITGB1 transfected cells**. D2A1 cells were transfected with control, anti-CD44 or anti-ITGB1 siRNAs. 72h later, cells were detached and perfused into microfluidic channels. The movie shows in parallel the perfusion of the three conditions at 100 μm/s. The movie is acquired at 24 fps and displayed at 72 fps. Related to Figure 2D.

**Movie 3 – Live adhesion movie of siCTL and siITGB3 transfected cells**. D2A1 cells were transfected with control or anti-ITGB3 siRNAs. 72h later, cells were detached and perfused into microfluidic channels. The movie shows in parallel the perfusion of the two conditions at 100 μm/s. The movie is acquired at 24 fps and displayed at 72 fps. Related to Figure 2D.

**Movie 4 – CD44 is not involved in stable adhesion *in vitro***. D2A1 cells were transfected with control siRNAs or anti-CD44. 72h later, cells were detached and perfused into microfluidic channels containing a confluent monolayer of endothelial cells (HUVEC) and left to attach to the endothelial layer without flow for 5 min. Attached cells were then trapped into the optical tweezer beam and the stage was moved away using the piezo stage. The movie is acquired at 24 fps and displayed at 72 fps. Related to Figure 2F.

**Movie 5 – ITGB3 is involved in stable adhesion *in vitro***. D2A1 cells were transfected with control siRNAs or anti-ITGB3. 72h later, cells were detached and perfused into microfluidic channels containing a confluent monolayer of endothelial cells (HUVEC) and left to attach to the endothelial layer without flow for 5 min. Attached cells were then trapped into the optical tweezer beam and the stage was moved away using the piezo stage. The movie was acquired at 24 fps and displayed at 72 fps. Related to Figure 2F.

**Movie 6 – ITGB1 is involved in stable adhesion *in vitro***. D2A1 cells were transfected with control siRNAs or anti-ITGB1. 72h later, cells were detached and perfused into microfluidic channels containing a confluent monolayer of endothelial cells (HUVEC) and left to attach to the endothelial layer without flow for 5 min. Attached cells were then trapped into the optical tweezer beam and the stage was moved away using the piezo stage. The movie was acquired at 24 fps and displayed at 72 fps. Related to Figure 2F.

**Movie 7 – CD44 is involved in CTC arrest/early adhesion *in vivo***. D2A1 cells were transfected with control or anti-CD44 siRNAs. 72h later, cells were detached and microinjected into the duct of Cuvier of 2 dpf *Tg(Fli1:EGFP)* embryos. Cell arrests was live imaged at 4 fps for 2,5 min immediately after injection. The movie is displayed at 80 fps. Related to Figure 3A-D.

**Movie 8 – ITGB3 is involved in CTC arrest/early adhesion *in vivo***. D2A1 cells were transfected with control or anti-ITGB1 siRNAs. 72h later, cells were detached and microinjected into the duct of Cuvier of 2 dpf *Tg(Fli1:EGFP)* embryos. Cell arrests was live imaged at 4 fps for 2,5 min immediately after injection. The movie is displayed at 80 fps. Related to Figure 3A-D.

**Movie 9 – ITGB1 is involved in CTC arrest/early adhesion *in vivo***. D2A1 cells were transfected with control or anti-ITGB1 siRNAs. 72h later, cells were detached and microinjected into the duct of Cuvier of 2 dpf *Tg(Fli1:EGFP)* embryos. Cell arrests was live imaged at 4 fps for 2,5 min immediately after injection. The movie is displayed at 80 fps. Related to Figure 3A-D.

